# Multi-curve fitting and tubulin-lattice signal removal for structure determination of large microtubule-based motors

**DOI:** 10.1101/2022.01.22.477366

**Authors:** Pengxin Chai, Qinhui Rao, Kai Zhang

## Abstract

Revealing high-resolution structures of microtubule-associated proteins (MAPs) is critical for understanding their fundamental roles in various cellular activities, such as cell motility and intracellular cargo transport. Nevertheless, large molecular motors that dynamically bind and release microtubule networks are challenging for cryo-electron microscopy (cryo-EM). Traditional structure determination of MAPs bound to microtubules needs alignment information from the reconstruction of microtubules, which cannot be readily applied to large MAPs without a fixed binding pattern. Here, we developed a comprehensive approach to estimate the microtubule networks (multi-curve fitting), model the tubulin-lattice signals, and remove them (tubulin-lattice subtraction) from the raw cryo-EM micrographs. The approach does not require an ordered binding pattern of MAPs on microtubules, nor does it need a reconstruction of the microtubules. We demonstrated the capability of our approach using the reconstituted outer-arm dynein bound to microtubule doublets. In addition, we applied our multi-curve fitting approach to other biological filaments and achieved accurate estimations. Our work provides a new tool to determine high-resolution structures of large MAPs bound to curved microtubule networks.

## 1. Introduction

The cryo-electron microscopy (cryo-EM) field has seen rapid development in the past few years *(1, 2)*. High-resolution structure determination of rigid globular proteins and helical assemblies in solution has become readily achievable in general cases *(3–6)*. However, there are still numerous challenges ahead, which require considerable efforts on careful sample preparation, optimization of data acquisition, or improved image processing approaches. One major challenge is to determine high-resolution structures of interconnected networks of cellular filaments decorated with intricate protein complexes, such as microtubules and the numerous microtubule-associated proteins (MAPs). Using *in vitro* reconstituted samples and improved image processing methods *(7, 8)*, high-resolution structures of several MAPs bound to microtubules have been obtained *(9–18)*. Moreover, a protofilament refinement protocol has been developed to further improve the reconstruction quality of each protofilament and their binding partners *(19)*.

Nevertheless, almost all currently reported structures of MAPs in the microtubule-binding states require a pre-reconstruction of the microtubule itself. Several key conditions must be met for a successful reconstruction in such cases. First, the MAPs must be small enough so that the microtubule alignment is not severely affected and a high-resolution reconstruction of the microtubule itself is achievable. Second, the decoration of the MAPs on microtubules must be specific, dense, and ordered to avoid insufficient signals of the targets after averaging. Finally, the bound MAPs must have stable conformations, sufficient orientations, and strong signals for convergent refinement. However, some MAPs, especially motor proteins, exhibit large conformational changes on microtubules. This leads to a common phenomenon that high-resolution features of MAPs can only be observed near the microtubule surface *(20–23)*.

Taking all these together, it is not straightforward to obtain high-resolution structures of large MAPs bound to microtubules in a less ordered or symmetry mismatched way using conventional single-particle approaches. Such limitation has been a major bottleneck that prevents us from understanding the detailed interactions between large motors and their tracks.

A well-established way to tackle symmetry mismatch and flexibility is to subtract the dominant signals from the raw particles after the three-dimensional (3D) reconstruction *(24–28)*. Similar approaches can be potentially applied to less ordered large MAPs. However, successful application of post-3D subtraction methods requires several key conditions, such as (i) accurate 3D reconstruction of the dominant signals, (ii) successful pass of alignment information to target signals, (iii) limited motion among different regions, and (iv) relatively clean background. For many intricate filamentous networks and their associated proteins, none of these conditions can be straightforwardly satisfied. New approaches are urgently needed to tackle those emerging challenges.

Here, we used the outer-arm dynein (OAD) from *Tetrahymena thermophila* as a good example to develop a comprehensive approach for high-resolution reconstruction of large, flexible, and symmetry-mismatched MAPs bound to the interwoven microtubule networks. OAD is a 1.8-MDa molecular motor that can spontaneously form an array with 24-nm periodicity upon microtubule doublet (MT doublet) binding *(29, 30)* (**Fig. 1a**). Nevertheless, multiple binding sites on the MT doublet surface co-exist during the *in-vitro* reconstitution assay *(31)*, which leads to mismatched symmetries among different segments. As a result, OAD signals appear completely smeared in the reconstructed MT doublet maps using conventional single-particle approaches (**Fig. 1b**). Furthermore, attempts to directly reconstruct OAD also fail because particle picking, 2D classification, and alignment are all severely influenced by the predominant signal from the tubulin lattice (**Fig. 1b**). On the other hand, as OAD is relatively large and multiple OAD units are densely arrayed on MT doublet, obtaining a reliable initial model of MT doublet itself is difficult possibly due to the severe signal interference from OAD molecules (**Fig. 1b**). Therefore, the conventional post-3D signal subtraction approach *(24, 26–28)* is not applicable for high-resolution analysis of less ordered large motors on microtubule networks.

**Fig. 1.**
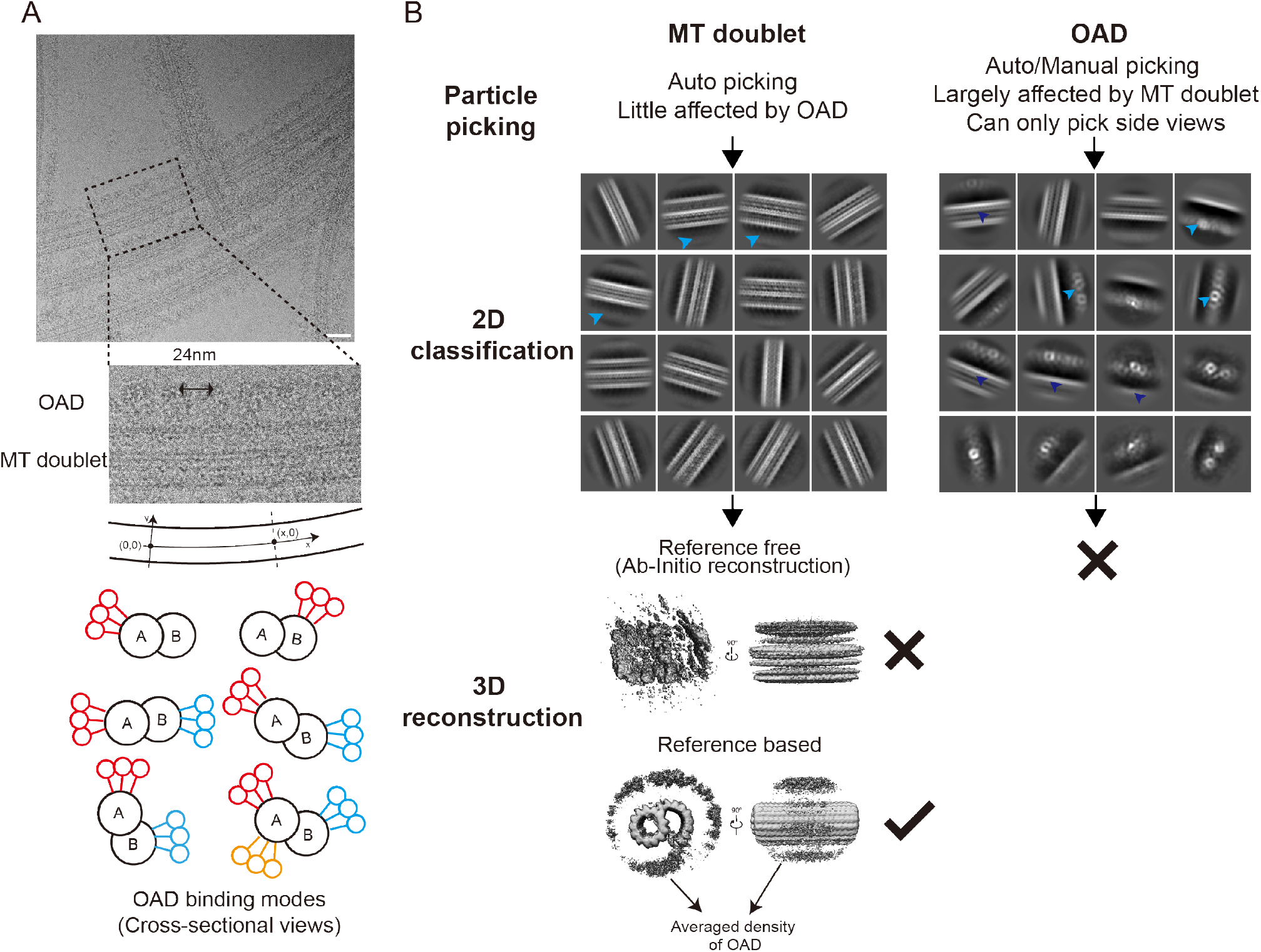
Outer-Arm Dynein (OAD) reconstruction is severely influenced by the signal from MT doublet. **(A)** Raw cryo-EM micrograph and schematic representation of OAD binding modes. The OAD and tubulin signal repeat along the major aixs of microtubule(x direction). Scale bar: 50 nm. **(B)** Cryo-EM image processing of MT doublet and OAD from the OAD-MT doublet reconstitution datasets. For MT doublet, the particle picking and 2D classification are little affected by the signal of OAD. 3D reconstruction of MT doublet is only possible given a reference. For OAD, every step of image processing is largely influenced by tubulin lattice signal. 3D reconstruction of OAD in the presence of MT doublet is not achievable. The OAD array and MT doublet are indicated by sky blue and blue arrows respectively.

Inspired by early studies on the statistical modeling and removal of liposome signals *(32, 33)*, we have developed a comprehensive approach to estimate microtubule networks, model the tubulin-lattice signals, and remove them from the raw micrographs, which allows accurate 3D reconstruction of OAD complexes in combination with a multiple-level local refinement procedure. The binding protofilaments signals and microtubule-OAD interactions can be accurately restored by subsequent local refinement of the original images using the alignment information from OAD. Our method fills an important gap of current approaches for cryo-EM studies on large MAPs and can be potentially applied to many other similar biological systems.

## 2. Material and methods

### 2.1 A general model for MAPs bound to microtubule networks

We aim to obtain high-resolution cryo-EM reconstructions of MAPs out of the datasets that meet the following challenging conditions: (i) highly curved and intersecting sets of microtubules, (ii) mismatched symmetries due to multiple binding sites, less ordered and sparse decoration of microtubules, (iii) large size and multi-level flexibility of the target MAPs. To solve these challenges, we developed an iterative approach to trace individual microtubules from the networks by fitting multiple polynomials simultaneously and removing the tubulin-lattice signals estimated from local 2D averages. The large MAPs can be subsequently treated as a normal singleparticle target and refined using a multi-level local refinement procedure to obtain high-resolution information of local regions.

### 2.2 Theoretical basis of tubulin-lattice signal modeling and subtraction

Microtubules are filamentous structures built by densely packed α/β-tubulin heterodimers that form individual protofilaments in a head-to-tail manner *(34)*. Depending on whether the path of each protofilament aligns with the major axis of the tubular structure, microtubules can be divided into straight or super-twisted classes *(35)*. The common 13-protofilament microtubule and MT doublet both belong to the straight class and have repeating tubulin subunits along the major axis per 8 nm. Considering the high structural similarity between the α- and β-tubulin *(36)*, tubulin-lattice signals repeat at approximately 4 nm intervals along the major axis. Meanwhile, large microtubule-based motor proteins may have different periodicities, such as OAD which repeats every 24 nm, or may just randomly bind to the microtubule, such as the dynein/dynactin/BICD2 complex.

Since the straight microtubule is a periodic 3D structure, the 2D projection of microtubule is also periodic. We can therefore treat the projection of each microtubule as an 1D crystal. When the MAP has a periodicity that differs from the microtubule, we can then perform signal decomposition by utilizing this symmetry discrepancy of the 1D crystal. After decomposition, the signal from microtubule can be subtracted without doing a 3D reconstruction.

A statistical model to deal with more general cases is simplified as follows. Suppose there exist two different types of signals *S*_1_(*x, y*) and *S*_2_(*x, y*), which repeat at periodicities *T*_1_ and *T*_2_ (*T*_2_>*T*_1_) along the *x* direction (*x*, curve length along the central major axis; *y*, distance to the central major axis). At any given position (*x, y*), the total signal with noise can be written as:

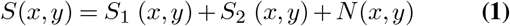

where *S*_1_ is the dominant signal with the smaller periodicity *T*_1_, while *S*_2_ is the signal of interest, of which the alignment is affected by *S*_1_; *N* is the noise. For simplicity, we ignore the background value here and will discuss it in detail later. In addition, since the periodic signals are restricted to the *x* direction in our model, we will ignore the *y*-axis in the following analyses:

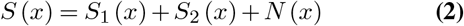

We estimate the averaged signal using *K* adjacent segments at *T*_1_ periodicity as follows:

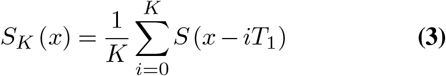

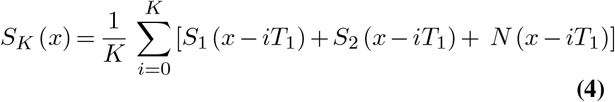

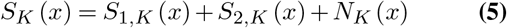

Here, the K-averaged value of signals at position x can be decomposed into three parts: (i) *S*_1,*K*_ (*x*), the contribution from the coherent average of the signal *S*_1_ at its own periodicity *T*_1_, (ii) *S*_2,*K*_ (*x*), the contribution from *S*_2_, which has a periodicity of *T*_2_, but is incoherently averaged using a smaller periodicity *T*_1_, (iii)*N_K_*, the noise after local average. Assume *N* ~ N(0,*σ*^2^), *N_K_* is also Gaussian and 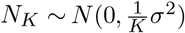.

Since *S*_1_ (*x*) = *S*_1_ (*x* − *iT*_1_), ∀*i* ∈ *Z*, the value of *S*_1,*K*_(*x*) is ideally *S*_1_(*x*) itself under the assumption that all other parameters are perfect. However, due to the imperfect estimation of *T*_1_ and positional parameters of each averaged segment, there is an image blurring effect which can be simply regarded as a convolution of *S*_1_ with a Gaussian kernel. This blurring effect does not significantly affect low-frequency information.

*S*_2,*K*_ (*x*) can be further decomposed into two parts, the contribution from *S*_2_ (*x*), equivalents at the position (*x* − *iT*_1_), where there exists an integer *j* (*i, j* ≤ *K*) such that *iT*_1_ = *jT*_2_, and signals from other non-equivalent positions. For typical biological filaments and their associated proteins, *T*_2_ is typically an integer multiple of *T*_1_ (e.g., 24-nm OAD repeat versus ‘4-nm’ MT doublet repeat). Assume *T*_2_ = *mT*_1_ and *K* is sufficiently big (e.g., >10m), *S*_2,*K*_ (*x*) can be approximate as:

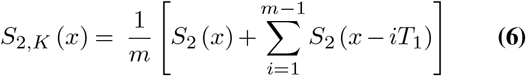

The background of cryo-EM images can vary significantly among different areas due to ice thickness or uneven distribution of electron beam intensity. However, the effect can be easily minimized by a high-pass filter *(37)* or real-space background estimation approaches *(33)*. Therefore, it can be reduced to a constant function as long as the images have been properly pre-processed, and its final effect can be ignored.

Now, we estimate the ideal information at x after subtracting *S_K_* from *S* based on the above assumptions and simplification.

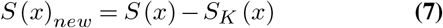

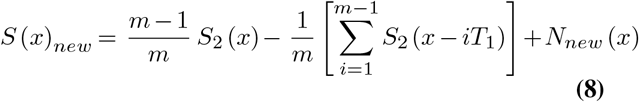

From the equation, it is obvious that the contribution from the signal *S*_1_ is gone in the ideal case. We linearly scale *S_new_* by a factor of 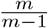:

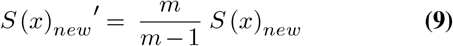

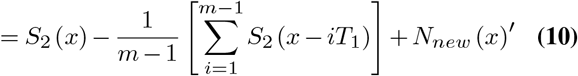

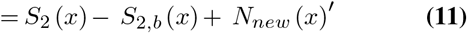

The information of the scaled image after *S_K_* subtraction contains three major parts: (i) the target signal *S*_2_, (ii) *S*_2,*b*_ (*x*) the bias imposed by averaging (*m* − 1) consecutively translated *S*_2_ signals, and (iii) the noise *N_new_′*, the new distribution of noise after subtraction, explicitly expressed as 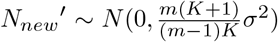. The noise *N_new_′* is slightly higher than the original noise *N* from the raw micrographs. When *K* and *m* are both sufficiently large, *N_new_′* is nearly the same as the original noise level. Thus, we conclude the additional noise imposed after subtraction is minor. The term *S*_2,*b*_ (*x*) can be regarded as a bias, which depends on the specific targets and may vary in different cases. The correlation between *S*_2_ and *S*_2,*b*_ reflects how much the target structure resembles itself by translation. In general cases, we assume *S*_2_ does not show any translational symmetries of its own, which will result in a significantly low correlation with *S*_2,*b*_, such that the normalized cross-correlation *NCC* (*S*_2_, *S*_2,*b*_) << 1, especially at high-frequency.

In the ideal case, the signal *S*_1_ is completely removed from the raw micrographs, while the target signal *S*_2_ is largely preserved except a minor addition of the bias *S*_2,*b*_ and slightly increased noise *N_new_′*. In practice, other factors also affect the accuracy of *S_new_′*, such as background estimation, curve fitting, periodicity, pseudo-symmetry (e.g., α- and β-tubulin), CTF variation, magnification distortion, structural flexibility, and other unexpected signals. These factors mostly affect the accuracy of high-frequency information, as the effects on low-frequency information can be easily approximated. A more accurate estimation of the final subtracted image is described as below:

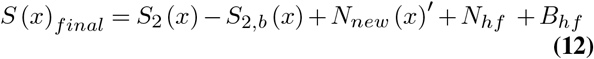

where *N_hf_* and *B_hf_* are respectively the high-frequency noise and bias imposed by the aforementioned factors. They will not significantly affect the reconstruction of *S*_2_ as the alignment is dominated by low-frequency signals.

In the case of sparse and random decoration, such as the microtubules associated with dynein/dynactin/BICD2 complexes, *T*_2_ and *m* are regarded as infinitely large, such that the term *S*_2,*b*_ is near zero. Therefore, the bias is not stronger than that in the case of a finite *T*_2_.

Taking MT doublet projection signal as *S*_1_ and its repeating length 4 nm as *T*_1_, and OAD projection signal as *S*_2_ and its repeating length 24 nm as *T*_2_, the workflow for the local 2D averaging and subtraction of MT doublet signal is shown in **Fig. 2**. Here the filaments are not completely straight, so the averaging process needs to be modified to take curvatures into account. The curvature is parameterized through a process we call multi-curve fitting, which is described in the next section. During the averaging step, a soft-edged rectangle mask is applied for each microtubule to avoid numerical artifacts (**Fig. 2a and b**). The background is estimated using the data values outside the mask (**Fig. 2b**). The width of this mask is dynamically changed according to the projected microtubule width (**Fig. 2c**). **Fig. 2d** shows a typical image after tubulin-lattice signal weakening. More details about the parameters and usage are in our code documentation (https://github.com/PengxinChai).

**Fig. 2.**
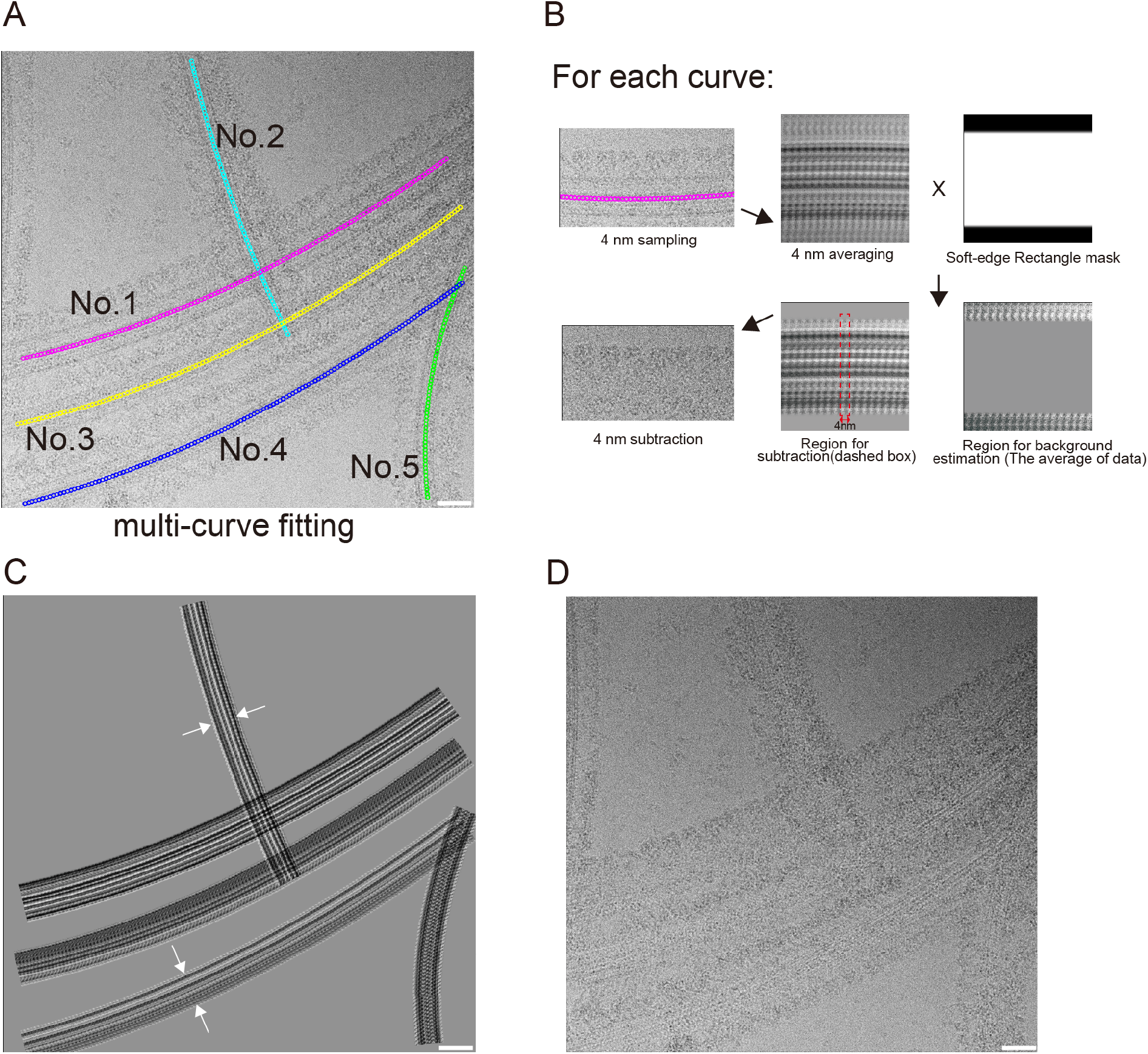
Tubulin-lattice signal modeling and subtraction from raw cryo-EM micrograph. **(A)** Multi-curve fitting of MT doublet filaments. **(B)** 4-nm tubulin-lattice signal modeling and subtraction. The major line of MT doublet was sampled at a 4 nm interval. Each 4 nm segment was extracted and aligned to generate an averaged image. A soft-edge rectangle mask was then applied. The center region was used for subtraction. The background value was estimated using the data points outside the mask. Finally, the subtraction between the original segment and 4 nm averaging results was performed. **(C)** The signal to be subtracted at the micrograph level. The white arrows indicated that the filament width was dynamically determined during the subtraction process. **(D)** The cryo-EM image after tubulin-lattice signal subtraction. Scale bar: 50 nm.

### 2.3 Multi-curve fitting for tracing microtubules

To obtain a high-quality estimate of *S_K_*(*x*) in the averaging process, the major axis of each filament needs to be accurately traced with the sampling points well centered and evenly spaced. We developed an iterative approach to fit and refine multiple curves out of the filament networks from cryo-EM micrographs (**Fig. 3a and b**). The flowchart contains four main steps, (a) initial sampling, (b) coordinate optimization, (c) multi-curve fitting, and (d) resampling. Using the reconstituted MT doublet-OAD as a typical target, more details of these steps are explained below.

**Fig. 3.**
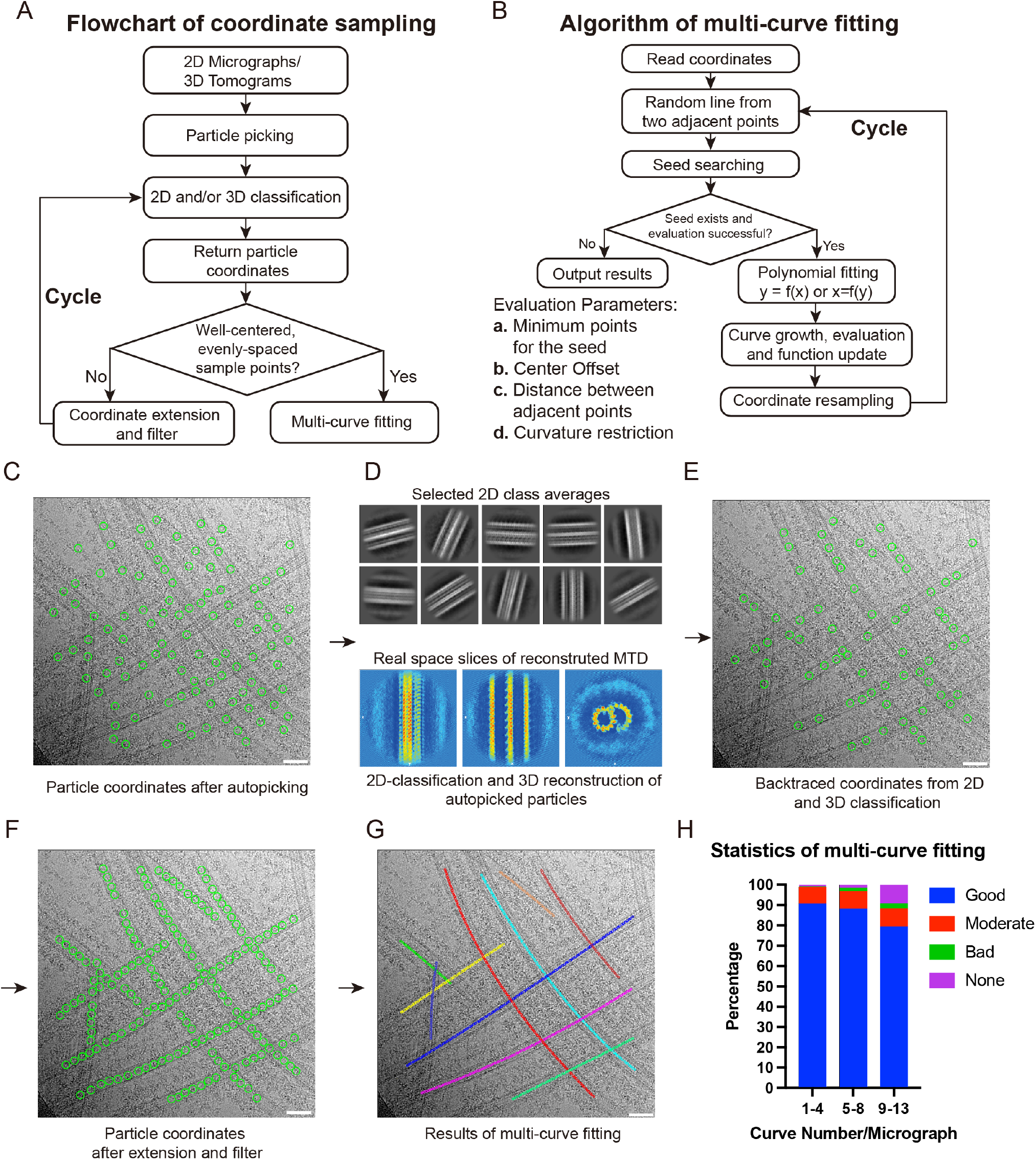
Coordinate sampling pipeline and multi-curve fitting algorithm. **(A)** Flowchart of coordinate sampling. **(B)** Flowchart of multi-curve fitting. **(C-F)** Representative results of coordinate sampling. **(G)** Representative result of multi-curve fitting. Images with coordinates were visualized using IMOD *(47)*. Scale bar: 50nm. **(H)** Statistical analysis of multi-curve fitting. Good fitting means the observed MT doublet filament matches well to the fitted curve. Moderate fitting applies to the situation where one MT doublet filament is cut into several segments. Bad fitting means the curve doesn’t overlap with the centerline of the filament. None fitting means the filament is not fitted with any curve. 1-4, n=162; 5-8, n=790; 9-13, n=368. See Figure. S1 for examples.

#### (a) Initial sampling

The goal is to obtain an initial set of coordinates as a rough estimate of microtubule segment locations. This can be done by automatic particle selection with or without templates using Gautomatch or other available programs *(38–40)*. Instead of trying to accurately detect all possible microtubule segments at the very beginning, our primary aim at this step is to coarsely assign sufficient seeding coordinates for subsequent optimization. Therefore, false positives can be largely tolerated during the initial sampling step. In addition, the particles along the filament do not need to be densely sampled (**Fig. 3c**).

#### (b) Coordinate optimization

The goal is to filter redundant or falsely picked particles, add missing ones, center all effective coordinates on the microtubules, and extract useful orientational information such as the in-plane rotation, which will minimize the errors due to insufficient and inaccurate sampling points for subsequent curve fitting. To achieve the goal, we perform 2D classification in Relion *(41)* or cryoSPARC *(42)* by combining the microtubule segments from all available cryo-EM datasets under the same condition, which allows improved signal-noise-ratios (SNR) and accuracy of particle centers after averaging. We subsequently extract the origin offsets of individual microtubule segments and re-center all of them. Falsely picked or low-quality particles are removed from the dataset through cycles of 2D classification. 3D reconstruction of the microtubule is not required at this step but can further improve the accuracy of particle centers and facilitate subsequent coordinate extension if feasible.

We add missing particles by three independent approaches, (i) interpolation of new coordinates between adjacent particles depending on their distances (does not need orientation information), (ii) coordinate extension towards both directions of each microtubule segment (need the in-plane rotation angle obtained from 2D classification or 3D reconstruction), (iii) extrapolation of the two ends of a certain filament after curve fitting. The three approaches can be combined in practice. Redundant coordinates are removed by a certain distance cutoff (e.g., <8 nm) (**Fig. 3d-f**).

#### (c) Multi-curve fitting

Due to the complexity of real-world problems, there is no single and simple solution to multi-curve fitting in general cases *(43, 44)*. We proposed the following approach to specifically optimize the multi-curve fitting problem of the microtubule networks from typical cryo-EM micrographs (**Fig. 3b**). The approach requires the optimized coordinates from step (b) as the input, under the assumption that our targets meet several empirical conditions, such as relatively small local curvatures (e.g., *k* < 1 *μm*^−1^), local smoothness, limited curve lengths, and a small number of crossovers from each micrograph. A summary of our approach is described below.

##### (c1) Seeding by line segments

A small subset of neighboring data points (typically ~ 5) is randomly assigned. Based on the assumption that the local curvature is small, a line segment is locally fitted by the least square minimization. Due to the random assignment, the local data points may come from single or multiple curves. If they belong to multiple microtubules, the line fitting error is normally extremely large and can be used as a criterion to reject the currently failed attempt. Once failed, new locations are attempted, while the failed points are marked to avoid reattempt.

##### (c2) Curve growth and evaluation

Once a seed of the local line segment is successfully found, it will start growing by absorbing the nearest neighbors from both ends. Distances and fitting errors between the absorbed points and the current seed are used as criteria to accept or reject the seed growth. Subsequently, a low-order (e.g., 2-3) polynomial function is locally fitted by the least square minimization on every successful seed. Each fitted curve then grows at both ends by taking in the neighbors that meet the cutoff criteria.

Two additional factors are considered to minimize the fitting errors. First, each polynomial function gets updated during the growth. Also, the abscissa and ordinate are swapped to make sure the deviation of the Y-coordinates from each micrograph is smaller than that of the X-coordinates, which is particularly helpful when a microtubule is nearly vertical in the cryo-EM micrograph.

##### (c3) Curve merging and re-assignment

For long microtubules, multiple seeds can likely be generated at different regions along with multiple polynomials fitted on a single filament. Two curves are merged into one if the difference between them is small and no significant gaps among the data points are detected. To reduce the possibility that some data points at the crossovers between two or more filaments are inaccurately assigned, we reassign each data point to the curve that results in a minimum fitting error. If there exists a significant subset of points (e.g., >5) that do not match any of the fitted curves. A new seed is generated out of those points for another cycle of multi-curve fitting and evaluation. Otherwise, the points that lead to large fitting errors are discarded. In practice, we manually check those abnormal micrographs.

#### (d) Resampling

With a polynomial function fitted, it is straightforward to generate a new set of well-centered and evenly spaced data points on each microtubule. We use the resampled coordinates to perform another cycle of 2D classification and/or 3D refinement. The optimized coordinates can be further utilized to re-perform the multi-curve fitting. In principle, the whole process can be iterated multiple times to potentially improve the accuracy of microtubule signal subtraction as described in Section 2.2. However, in practice, one additional cycle is sufficient, and more cycles do not significantly improve.

## 3. Results

### 3.1 Multi-curve fitting of microtubule networks

A representative result is shown in **Fig. 3c-g**. To evaluate the fitting accuracy of our methods, we randomly selected 200 micrographs and divided them into three groups based on the filament number. Then we compared the fitted curves and the actual filaments and assigned each of the filaments into one of the four categories: good, moderate, bad, and none fitted (**Fig. 3h and Figure S1**). Overall, the fitting accuracy of our approach is affected by the number of microtubules per micrograph. When the filament number per micrograph is less than 9, the result is sufficiently accurate (>89%). When it is over 10, the number of non-fitted and falsely fitted filaments increases (**Fig. 3h**). Among all the bad or non-fitting cases, only 1.2% of the filaments belong to the micrographs that contain no more than 5 microtubules each.

### 3.2 Tubulin-lattice signal subtraction substantially improves the alignment of OAD

The alignment of OAD after tubulin-lattice subtraction was substantially improved, as demonstrated by the new 2D class averages of OAD (**Fig. 4a**). The approach also allowed us to observe high-quality 2D class averages of residual radial spokes (RS) even if the most had been washed away by high salt treatment during MT doublet purification (**Fig. 4a**). Compared to the 2D class averages from original micrographs (**Fig. 1b**), the new results clearly show the features of dynein, such as the motor domain, tail region, and stalks. Most importantly, the top views of OAD were observed, suggesting that our approach can specifically weaken the tubulin-lattice signals and preserve the dynein signals. After extensive 2D and 3D classification, the final 288, 224 high-quality particles were used for 3D reconstruction, which yielded an overall 8.96 Å resolution map of OAD array bound to four protofilaments after global refinement (**Fig. 4b**). The resolution of the alpha-motor domain was improved to 3.19 Å after multi-level focused refinement (**Fig. 4b**) *(31)*. By contrast, no reliable 3D reconstruction could be achieved without multi-curve fitting and tubulin-lattice subtraction.

**Fig. 4.**
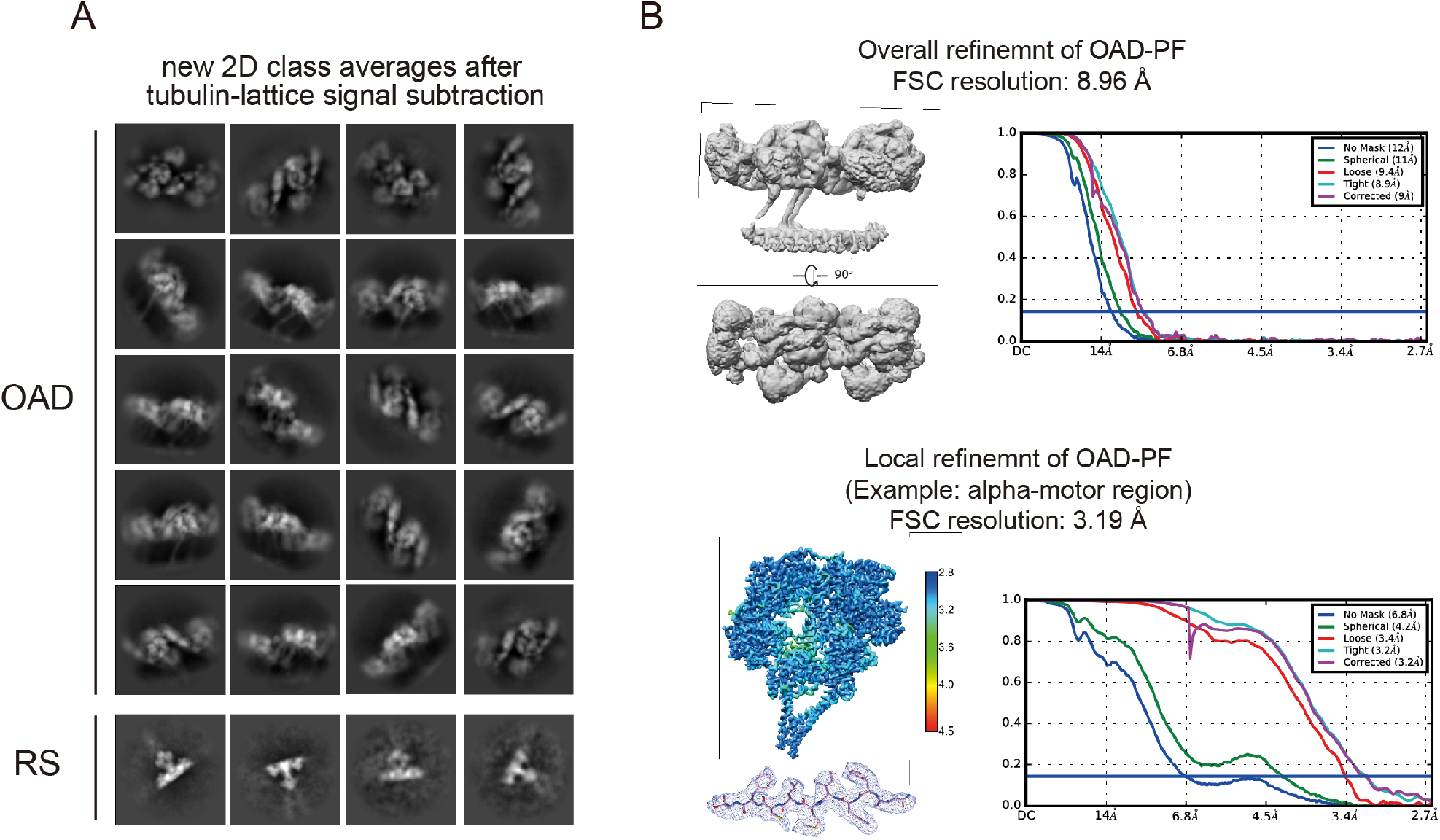
OAD reconstruction after tubulin-lattice signal subtraction. **(A)** Selected 2D class averages of OAD and radial spoke (RS). The averages clearly showed the features of dynein such as stalk, tail domain, and motor ring. **(B)** 3D reconstruction of the entire OAD bound to four protofilaments yielded an 8.96 Å resolution map. Subsequent local refinement on the alpha-motor region improved the resolution to 3.19 Å. Fourier Shell Correlation (FSC) curves were generated in Cryosparc *(42)*.

### 3.3 Application in other filamentous structures and 3D tomograms

Despite the original purpose to deal with the microtubule-based motors, our methods can be generalized to trace other biological filaments in both 2D micrographs and 3D tomograms. Singlet microtubules are more common than doublet microtubules during *in-vitro* reconstitution experiments, and our results demonstrate that our approaches could trace every single microtubule at high accuracy, including the challenging cases (**Figure S2A**). We also tested another filament, the cytochrome OmcS nanowire *(45)*. OmcS monomer from *G. sulfurreducens* polymerizes to form OmcS nanowires for electron conductivity. We started with automatic particle picking with a low cross-correlation threshold to have a good initial sampling of filaments despite some false positive particles. After 3 rounds of coordinates extension, filtering, and classification, nearly all the filament segments were well-centered and evenly spaced (**Figure S2B**). Our multi-curve fitting result shows that those filaments are accurately traced.

We also tested our multi-curve fitting algorithm in the case of 3D tomograms of the axoneme from *T. thermophila*. The seeding coordinates of MT doublet segments were first generated by template-based 3D particle picking in EMAN2 *(46)*. To extend our approach to 3D, we decompose the 3D coordinates into two sets of 2D coordinates from each tomogram, in the X-Y and X-Z planes respectively, and then performed multi-curve fitting separately as described in the case of 2D. We then compose a curve in 3D space from the two fitted curves in 2D. Finally, we evenly resample the (X, Y, Z) coordinates in 3D space using the fitted curve. The fitting results in 3D are sufficiently accurate (**Figure S2C**), which allows precise localization of the axonemal components such as the outer-arm dynein.

## 4. Discussion and conclusions

Microtubule reconstruction-based approaches have been developed and used to obtain several high-resolution structures of MAPs. This requires that MAPs be small or have defined binding sites such as proteins natively bound to MT doublet *(21, 22, 48)* to avoid the problem of symmetry mismatch. Our study provides a new tool to study the high-resolution structures of large MAPs bound to the microtubule with any possible pattern, no matter it is dense, sparse, ordered, or random, as long as the periodicity is not identical to that of the tubulins. The curvature estimation and local averaging of filaments in 2D avoids the difficulty of 3D modeling of the helical (or near-helical) structure. In the case of the reconstituted OAD arrays bound to MT doublets, we can also observe 2D class averages of radial spokes after tubulin-lattice subtraction (**Fig. 4**), suggesting that our method can be used for other protein complexes, especially for large proteins or protein complexes having a less ordered or sparse decoration on microtubules. In addition to microtubule surface-binding proteins, our approach can potentially be applied to the proteins in the microtubule lumen, such as recently published MAP6 that binds to the microtubule lumen at a periodicity of 31 nm *(49)* and actin filaments inside the microtubule lumen *(50)*. In addition, our approach provides a promising solution to high-resolution structure determination of dynein-dynactin-adaptor complexes bound to microtubules. Moreover, the same principle applies to many other biological systems other than microtubules, such as the host factors bound to viral capsid tubes.

In our study, we also developed a tool to iteratively trace filamentous objects by assembling several standard steps of signal-particle cryo-EM data processing. This is different from previously described filament tracing methods that attempt to segment filaments in the first place *(39, 40)*. We find the goal is normally infeasible in practice due to many unexpected factors that may affect the results. By contrast, our solution essentially regards the filaments as normal single particles and avoids extensive efforts to optimize the tracing parameters at the very beginning, but resorts to the information accumulated in the later steps and refines the tracing iteratively. After an initial selection of seeding particles, the positional and qualitative information of those particles can be optimized over cycles of iteration, which allows more neighboring particles of high quality to be identified. Theoretically, for a straight filament, one particle along the filament followed by one round of refinement and coordinate extension is sufficient to trace the entire filament. In the case of severely curved and overlapped filaments, the goal is more challenging but can be mitigated by more cycles of multi-curve fitting, classification, and particle optimization as described in Section 2.2.

We have demonstrated that the capability of our approach using the microtubule networks and bacterial nanowire OmcS and expect to expand the approach to other filamentous objects. For special filaments whose curvature cannot be approximated by polynomial functions, further optimization is needed to make the algorithm applicable. For example, a possible solution is to divide a very complicated curve into several short segments with sufficient overlaps, perform the multi-curve fitting and coordinate extension on these short curves, and finally smoothly ‘stitch’ them together.

## Acknowledgements

We thank F. Sigworth, C. Sindelar and Y. Xiong for valuable discussions, and N. Malvankar for providing the OmcS nanowire dataset.

## Funding and additional information

This work was supported by start-up funds from Yale University, the National Institutes of Health grants R35GM142959 awarded to K.Z. and S10OD023603 awarded to F. Sigworth, Rudolf J. Anderson Fellowship awarded to Q.R., and the China Scholarship Council (CSC)-Yale World Scholars Program in Biomedical Sciences awarded to P.C.

## Author contributions

K.Z. designed the project. P.C. wrote the program under K.Z.’s guidance. Q.R., P.C. and K.Z. tested the method, analyzed the OAD datasets and determined the structures. K.Z. and P.C. prepared the manuscript.

## Code availability

Source code and documentation is available at https://github.com/PengxinChai

---

**Figure S1.**
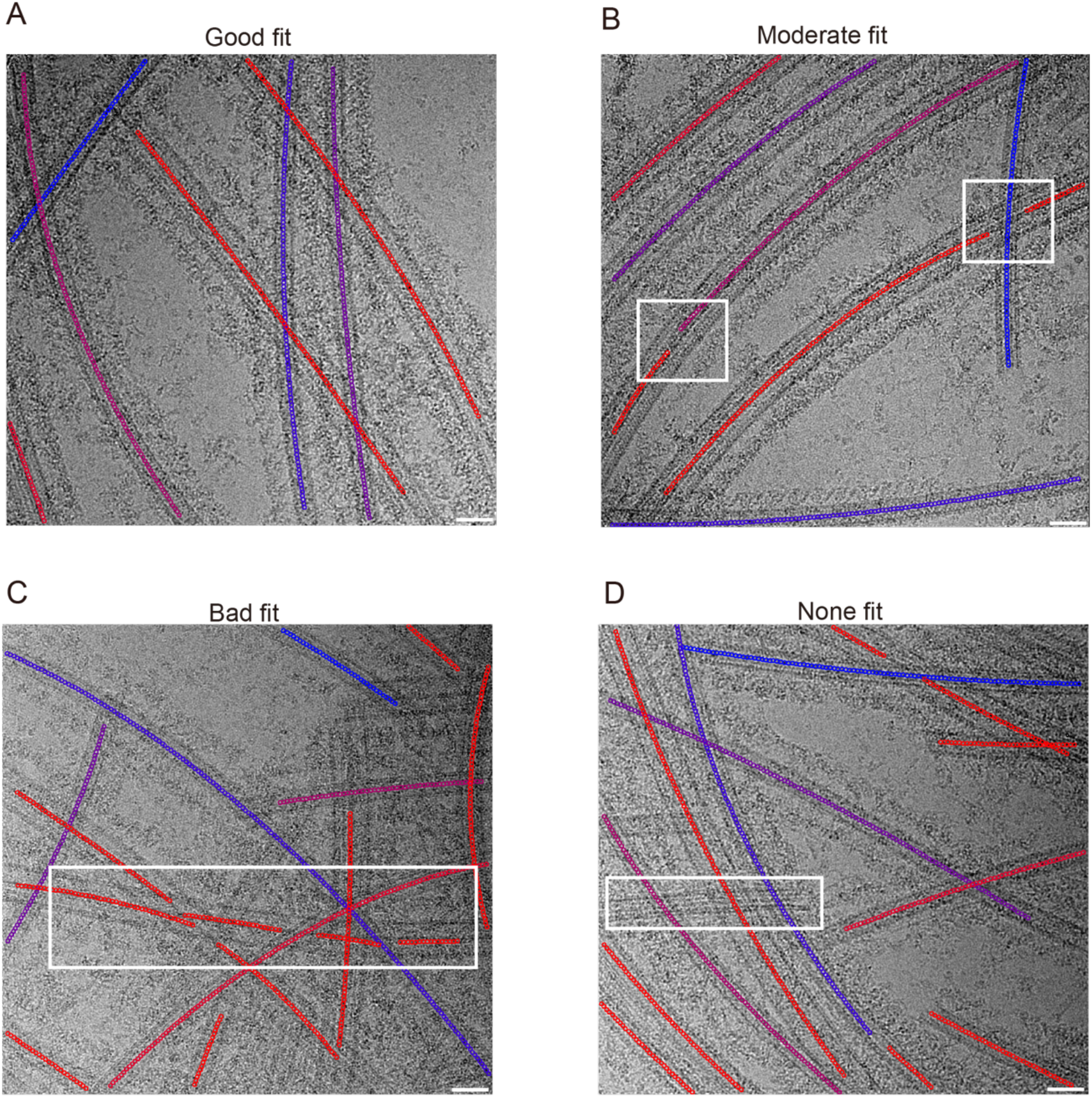
Representative results of multi-curve fitting in different situations. A: Good fit. The fitted curve traces the centerline of MT doublet well. B: Moderate fit shown in white rectangle boxes. Due to the crossover of filaments or the curvature, some filaments are fitted with more than one curve. C: Bad fit shown in a white rectangle box. The fitted curve doesn’t trace the centerline of the filament due to multiple crossovers. D: White rectangle box shows that some filaments are not fitted due to a combination effect from crossover and short length of filaments.

**Figure S2.**
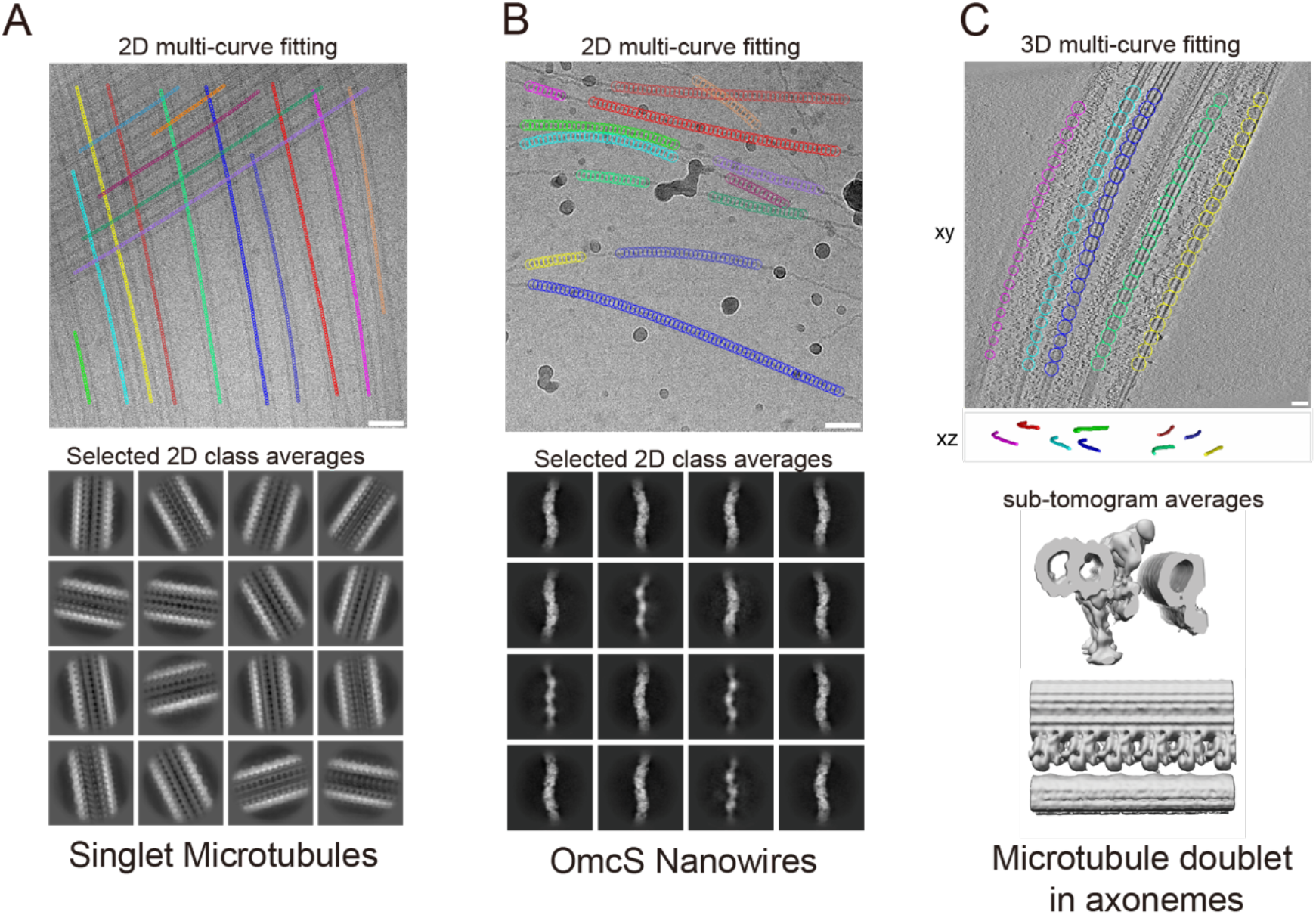
Application of coordinate sampling and multi-curve fitting. A: Representative results for microtubule singlets. Microtubule singlets dataset is downloaded from EMPIAR (EMPIAR-10300). B: Representative results for OmcS nanowire. C: 3D multi-curve fitting for the densely packed MT doublets in axonemes. After curve fitting, particle positions along with two Euler angles were extracted and imposed during subsequent sub-tomogram averaging analysis.

## References

1. Y. Cheng, N. Grigorieff, P. A. Penczek, and T. Walz. A primer to singleparticle cryo-electron microscopy. Cell, 161(3):438–449, 2015. ISSN 1097-4172 (Electronic) 0092-8674 (Linking). doi: 10.1016/j.cell.2015.03.050.

2. K. Murata and M. Wolf. Cryo-electron microscopy for structural analysis of dynamic biological macromolecules. Biochim Biophys Acta Gen Subj, 1862 (2):324–334, 2018. ISSN 0304-4165 (Print) 0304-4165 (Linking). doi: 10.1016/j.bbagen.2017.07.020.

3. A. Merk, A. Bartesaghi, S. Banerjee, V. Falconieri, P. Rao, M. I. Davis, R. Pragani, M. B. Boxer, L. A. Earl, J. L. S. Milne, and S. Subramaniam. Breaking cryo-em resolution barriers to facilitate drug discovery. Cell, 165(7): 1698–1707, 2016. ISSN 1097-4172 (Electronic) 0092-8674 (Linking). doi: 10.1016/j.cell.2016.05.040.

4. A. Bartesaghi, C. Aguerrebere, V. Falconieri, S. Banerjee, L. A. Earl, X. Zhu, N. Grigorieff, J. L. S. Milne, G. Sapiro, X. Wu, and S. Subramaniam. Atomic resolution cryo-em structure of beta-galactosidase. Structure, 26(6):848–856 e3, 2018. ISSN 1878-4186 (Electronic) 0969-2126 (Linking). doi: 10.1016/j.str.2018.04.004.

5. A. K. Mitra. Visualization of biological macromolecules at near-atomic resolution: cryo-electron microscopy comes of age. Acta Crystallographica Section F-Structural Biology Communications, 75:3–11, 2019. doi: 10.1107/S2053230x18015133.

6. S. He and S. H. W. Scheres. Helical reconstruction in relion. J Struct Biol, 198 (3):163–176, 2017. ISSN 1095-8657 (Electronic) 1047-8477 (Linking). doi: 10.1016/j.jsb.2017.02.003.

7. R. Zhang and E. Nogales. A new protocol to accurately determine microtubule lattice seam location. J Struct Biol, 192(2):245–54, 2015. ISSN 1095-8657 (Electronic) 1047-8477 (Linking). doi: 10.1016/j.jsb.2015.09.015.

8. A. D. Cook, S. W. Manka, S. Wang, C. A. Moores, and J. Atherton. A microtubule relion-based pipeline for cryo-em image processing. J Struct Biol, 209(1):107402, 2020. ISSN 1095-8657 (Electronic) 1047-8477 (Linking). doi: 10.1016/j.jsb.2019.10.004.

9. R. Zhang, G. M. Alushin, A. Brown, and E. Nogales. Mechanistic origin of microtubule dynamic instability and its modulation by eb proteins. Cell, 162 (4):849–59, 2015. ISSN 1097-4172 (Electronic) 0092-8674 (Linking). doi: 10.1016/j.cell.2015.07.012.

10. E. H. Kellogg, S. Howes, S. C. Ti, E. Ramirez-Aportela, T. M. Kapoor, P. Chacon, and E. Nogales. Near-atomic cryo-em structure of prc1 bound to the microtubule. Proc Natl Acad Sci U S A, 113(34):9430–9, 2016. ISSN 1091-6490 (Electronic) 0027-8424 (Linking). doi: 10.1073/pnas.1609903113.

11. D. Liu, X. Liu, Z. Shang, and C. V. Sindelar. Structural basis of cooperativity in kinesin revealed by 3d reconstruction of a two-head-bound state on microtubules. Elife, 6, 2017. ISSN 2050-084X (Electronic) 2050-084X (Linking). doi: 10.7554/eLife.24490.

12. R. Zhang, J. Roostalu, T. Surrey, and E. Nogales. Structural insight into tpx2-stimulated microtubule assembly. Elife, 6, 2017. ISSN 2050-084X (Electronic) 2050-084X (Linking). doi: 10.7554/eLife.30959.

13. Mpmh Benoit, A. B. Asenjo, and H. Sosa. Cryo-em reveals the structural basis of microtubule depolymerization by kinesin-13s. Nat Commun, 9(1): 1662, 2018. ISSN 2041-1723 (Electronic) 2041-1723 (Linking). doi: 10.1038/s41467-018-04044-8.

14. E. H. Kellogg, N. M. A. Hejab, S. Poepsel, K. H. Downing, F. DiMaio, and E. Nogales. Near-atomic model of microtubule-tau interactions. Science, 360 (6394):1242–1246, 2018. ISSN 1095-9203 (Electronic) 0036-8075 (Linking). doi: 10.1126/science.aat1780.

15. F. Li, Y. Li, X. Ye, H. Gao, Z. Shi, X. Luo, L. M. Rice, and H. Yu. Cryo-em structure of vash1-svbp bound to microtubules. Elife, 9, 2020. ISSN 2050-084X (Electronic) 2050-084X (Linking). doi: 10.7554/eLife.58157.

16. A. Pena, A. Sweeney, A. D. Cook, J. Locke, M. Topf, and C. A. Moores. Structure of microtubule-trapped human kinesin-5 and its mechanism of inhibition revealed using cryoelectron microscopy. Structure, 28(4):450–457 e5, 2020. ISSN 1878-4186 (Electronic) 0969-2126 (Linking). doi: 10.1016/j.str.2020.01.013.

17. T. Bodrug, E. M. Wilson-Kubalek, S. Nithianantham, A. F. Thompson, A. Alfieri, I. Gaska, J. Major, G. Debs, S. Inagaki, P. Gutierrez, L. Gheber, R. J. McKenney, C. V. Sindelar, R. Milligan, J. Stumpff, S. S. Rosenfeld, S. T. Forth, and J. Al-Bassam. The kinesin-5 tail domain directly modulates the mechanochemical cycle of the motor domain for anti-parallel microtubule sliding. Elife, 9, 2020. ISSN 2050-084X (Electronic) 2050-084X (Linking). doi: 10.7554/eLife.51131.

18. S. E. Lacey, S. He, S. H. Scheres, and A. P. Carter. Cryo-em of dynein microtubule-binding domains shows how an axonemal dynein distorts the microtubule. Elife, 8, 2019. ISSN 2050-084X (Electronic) 2050-084X (Linking). doi: 10.7554/eLife.47145.

19. G. E. Debs, M. Cha, X. Liu, A. R. Huehn, and C. V. Sindelar. Dynamic and asymmetric fluctuations in the microtubule wall captured by high-resolution cryoelectron microscopy. Proc Natl Acad Sci U S A, 117(29):16976–16984, 2020. ISSN 1091-6490 (Electronic) 0027-8424 (Linking). doi: 10.1073/pnas.2001546117.

20. M. Ma, M. Stoyanova, G. Rademacher, S. K. Dutcher, A. Brown, and R. Zhang. Structure of the decorated ciliary doublet microtubule. Cell, 179(4):909–922 e12, 2019. ISSN 1097-4172 (Electronic) 0092-8674 (Linking). doi: 10.1016/j.cell.2019.09.030.

21. M. Gui, M. Ma, E. Sze-Tu, X. Wang, F. Koh, E. D. Zhong, B. Berger, J. H. Davis, S. K. Dutcher, R. Zhang, and A. Brown. Structures of radial spokes and associated complexes important for ciliary motility. Nat Struct Mol Biol, 28(1):29–37, 2021. ISSN 1545-9985 (Electronic) 1545-9985 (Linking). doi: 10.1038/s41594-020-00530-0.

22. T. Walton, H. Wu, and A. Brown. Structure of a microtubule-bound axonemal dynein. Nat Commun, 12(1):477, 2021. ISSN 2041-1723 (Electronic) 2041-1723 (Linking). doi: 10.1038/s41467-020-20735-7.

23. S. Kubo, S. K. Yang, C. S. Black, D. Dai, M. Valente-Paterno, J. Gaertig, M. Ichikawa, and K. H. Bui. Remodeling and activation mechanisms of outer arm dyneins revealed by cryo-em. EMBO Rep, 22(9):e52911, 2021. ISSN 1469-3178 (Electronic) 1469-221X (Linking). doi: 10.15252/embr.202152911.

24. X. C. Bai, E. Rajendra, G. Yang, Y. Shi, and S. H. Scheres. Sampling the conformational space of the catalytic subunit of human gamma-secretase. Elife, 4, 2015. ISSN 2050-084X (Electronic) 2050-084X (Linking). doi: 10.7554/eLife.11182.

25. T. Nakane, D. Kimanius, E. Lindahl, and S. H. Scheres. Characterisation of molecular motions in cryo-em single-particle data by multi-body refinement in relion. Elife, 7, 2018. ISSN 2050-084X (Electronic) 2050-084X (Linking). doi: 10.7554/eLife.36861.

26. S. L. Ilca, A. Kotecha, X. Sun, M. M. Poranen, D. I. Stuart, and J. T. Huiskonen. Localized reconstruction of subunits from electron cryomicroscopy images of macromolecular complexes. Nat Commun, 6:8843, 2015. ISSN 2041-1723 (Electronic) 2041-1723 (Linking). doi: 10.1038/ncomms9843.

27. H. Liu and L. Cheng. Cryo-em shows the polymerase structures and a nonspooled genome within a dsrna virus. Science, 349(6254):1347–50, 2015. ISSN 1095-9203 (Electronic) 0036-8075 (Linking). doi: 10.1126/science.aaa4938.

28. J. T. Huiskonen, H. T. Jaalinoja, J. A. Briggs, S. D. Fuller, and S. J. Butcher. Structure of a hexameric rna packaging motor in a viral polymerase complex. J Struct Biol, 158(2):156–64, 2007. ISSN 1047-8477 (Print) 1047-8477 (Linking). doi: 10.1016/j.jsb.2006.08.021.

29. M. E. Porter and K. A. Johnson. Characterization of the atp-sensitive binding of tetrahymena 30 s dynein to bovine brain microtubules. J Biol Chem, 258 (10):6575–81, 1983. ISSN 0021-9258 (Print) 0021-9258 (Linking).

30. T. Oda, T. Abe, H. Yanagisawa, and M. Kikkawa. Docking-complex-independent alignment of chlamydomonas outer dynein arms with 24-nm periodicity in vitro. J Cell Sci, 129(8):1547–51, 2016. ISSN 1477-9137 (Electronic) 0021-9533 (Linking). doi: 10.1242/jcs.184598.

31. Q. Rao, L. Han, Y. Wang, P. Chai, Y. W. Kuo, R. Yang, F. Hu, Y. Yang, J. Howard, and K. Zhang. Structures of outer-arm dynein array on microtubule doublet reveal a motor coordination mechanism. Nat Struct Mol Biol.

32. K. H. Jensen, S. S. Brandt, H. Shigematsu, and F. J. Sigworth. Statistical modeling and removal of lipid membrane projections for cryo-em structure determination of reconstituted membrane proteins. J Struct Biol, 194(1):49–60, 2016. ISSN 1095-8657 (Electronic) 1047-8477 (Linking). doi: 10.1016/j.jsb.2016.01.012.

33. K. H. Jensen, F. J. Sigworth, and S. S. Brandt. Removal of vesicle structures from transmission electron microscope images. IEEE Trans Image Process, 25(2):540–52, 2016. ISSN 1941-0042 (Electronic) 1057-7149 (Linking). doi: 10.1109/TIP.2015.2504901.

34. A. Desai and T. J. Mitchison. Microtubule polymerization dynamics. Annu Rev Cell Dev Biol, 13:83–117, 1997. ISSN 1081-0706 (Print) 1081-0706 (Linking). doi: 10.1146/annurev.cellbio.13.1.83.

35. S. Chaaban and G. J. Brouhard. A microtubule bestiary: structural diversity in tubulin polymers. Mol Biol Cell, 28(22):2924–2931, 2017. ISSN 1939-4586 (Electronic) 1059-1524 (Linking). doi: 10.1091/mbc.E16-05-0271.

36. J. Lowe, H. Li, K. H. Downing, and E. Nogales. Refined structure of alpha beta-tubulin at 3.5 a resolution. J Mol Biol, 313(5):1045–57, 2001. ISSN 0022-2836 (Print) 0022-2836 (Linking). doi: 10.1006/jmbi.2001.5077.

37. X. Fan, J. Wang, X. Zhang, Z. Yang, J. C. Zhang, L. Zhao, H. L. Peng, J. Lei, and H. W. Wang. Single particle cryo-em reconstruction of 52 kda streptavidin at 3.2 angstrom resolution. Nat Commun, 10(1):2386, 2019. ISSN 2041-1723 (Electronic) 2041-1723 (Linking). doi: 10.1038/s41467-019-10368-w.

38. T. Wagner, F. Merino, M. Stabrin, T. Moriya, C. Antoni, A. Apelbaum, P. Hagel, O. Sitsel, T. Raisch, D. Prumbaum, D. Quentin, D. Roderer, S. Tacke, B. Siebolds, E. Schubert, T. R. Shaikh, P. Lill, C. Gatsogiannis, and S. Raunser. Sphire-cryolo is a fast and accurate fully automated particle picker for cryo-em. Commun Biol, 2:218, 2019. ISSN 2399-3642 (Electronic) 2399-3642 (Linking). doi: 10.1038/s42003-019-0437-z.

39. T. Wagner, L. Lusnig, S. Pospich, M. Stabrin, F. Schonfeld, and S. Raunser. Two particle-picking procedures for filamentous proteins: Sphire-cryolo filament mode and sphire-striper. Acta Crystallogr D Struct Biol, 76(Pt 7): 613–620, 2020. ISSN 2059-7983 (Electronic) 2059-7983 (Linking). doi: 10.1107/S2059798320007342.

40. S. T. Huber, T. Kuhm, and C. Sachse. Automated tracing of helical assemblies from electron cryo-micrographs. J Struct Biol, 202(1):1–12, 2018. ISSN 1095-8657 (Electronic) 1047-8477 (Linking). doi: 10.1016/j.jsb.2017.11.013.

41. J. Zivanov, T. Nakane, B. O. Forsberg, D. Kimanius, W. J. Hagen, E. Lindahl, and S. H. Scheres. New tools for automated high-resolution cryo-em structure determination in relion-3. Elife, 7, 2018. ISSN 2050-084X (Electronic) 2050-084X (Linking). doi: 10.7554/eLife.42166.

42. A. Punjani, J. L. Rubinstein, D. J. Fleet, and M. A. Brubaker. cryosparc: algorithms for rapid unsupervised cryo-em structure determination. Nat Methods, 14(3):290–296, 2017. ISSN 1548-7105 (Electronic) 1548-7091 (Linking). doi: 10.1038/nmeth.4169.

43. P. Spitzer, C. Zierhofer, and E. Hochmair. Algorithm for multi-curve-fitting with shared parameters and a possible application in evoked compound action potential measurements. Biomed Eng Online, 5:13, 2006. ISSN 1475-925X (Electronic) 1475-925X (Linking). doi: 10.1186/1475-925X-5-13.

44. Ieng S S. Tarel J P, Charbonnier P. Simultaneous robust fitting of multiple curves. VISAPP, 175–182, 2007.

45. F. B. Wang, Y. Q. Gu, J. P. O’Brien, S. M. Yi, S. E. Yalcin, V. Srikanth, C. Shen, D. Vu, N. L. Ing, A. I. Hochbaum, E. H. Egelman, and N. S. Malvankar. Structure of microbial nanowires reveals stacked hemes that transport electrons over micrometers. Cell, 177(2):361–+, 2019. ISSN 0092-8674. doi: 10.1016/j.cell.2019.03.029.

46. G. Tang, L. Peng, P. R. Baldwin, D. S. Mann, W. Jiang, I. Rees, and S. J. Ludtke. Eman2: an extensible image processing suite for electron microscopy. J Struct Biol, 157(1):38–46, 2007. ISSN 1047-8477 (Print) 1047-8477 (Linking). doi: 10.1016/j.jsb.2006.05.009.

47. J. R. Kremer, D. N. Mastronarde, and J. R. McIntosh. Computer visualization of three-dimensional image data using imod. J Struct Biol, 116(1):71–6, 1996. ISSN 1047-8477 (Print) 1047-8477 (Linking). doi: 10.1006/jsbi.1996.0013.

48. A. A. Z. Khalifa, M. Ichikawa, D. Dai, S. Kubo, C. S. Black, K. Peri, T. S. McAlear, S. Veyron, S. K. Yang, J. Vargas, S. Bechstedt, J. F. Trempe, and K. H. Bui. The inner junction complex of the cilia is an interaction hub that involves tubulin post-translational modifications. Elife, 9, 2020. ISSN 2050-084X (Electronic) 2050-084X (Linking). doi: 10.7554/eLife.52760.

49. C. Cuveillier, J. Delaroche, M. Seggio, S. Gory-Faure, C. Bosc, E. Denarier, M. Bacia, G. Schoehn, H. Mohrbach, I. Kulic, A. Andrieux, I. Arnal, and C. Delphin. Map6 is an intraluminal protein that induces neuronal microtubules to coil. Science Advances, 6(14), 2020. ISSN 2375-2548. doi: ARTNeaaz434410.1126/sciadv.aaz4344.

50. D. M. Paul, J. Mantell, U. Borucu, J. Coombs, K. J. Surridge, J. M. Squire, P. Verkade, and M. P. Dodding. In situ cryo-electron tomography reveals filamentous actin within the microtubule lumen. Journal of Cell Biology, 219(9), 2020. ISSN 0021-9525. doi: ARTNe20191115410.1083/jcb.201911154.

